# B-cell-Specific *Wwox* Deletion Promotes Plasmablastic Tumor Development and Pro-Inflammatory Signatures in a Myeloma Mouse Model

**DOI:** 10.1101/2025.01.29.635577

**Authors:** Tabish Hussain, Matthew D. Bramble, Bin Liu, Martin C. Abba, Marta Chesi, C. Marcelo Aldaz

## Abstract

Deletions and translocations affecting *WWOX* accompanied by loss of expression are frequently observed in B cell neoplasms and are linked to poor prognosis. Our previous research showed that *Wwox* deletion early in B cell development induces genomic instability, neoplastic transformation, and monoclonal gammopathies in mice. In this study, by crossing *Cd19 Wwox* knockout (KO) with *Vk*MYC* myeloma model mice, we generated a model with concurrent *Wwox* deletion and *MYC* activation, reproducing two common oncogenic alterations in B and plasma cell cancers. We observed that *Vk*MYC:Wwox KO* mice exhibited significantly reduced survival rates primarily due to the development of plasmablastic plasmacytomas and lymphomas. Transcriptome profiling from bone marrow derived Cd138+ plasma cells and plasmablastic tumors revealed enrichment of biofunctions related to tumorigenic phenotype and inflammation activation upon *Wwox* deletion in *Vk*MYC* mice. *Wwox KO* plasmablastic tumors displayed mutations affecting classical cancer genes, DNA damage response (DDR) genes, as well as overexpression of Aid/Apobec family members associated to hypermutation and DDR mutational signatures. These findings illustrate the significant pathobiological effects of B cell specific *Wwox* deletion and support a relevant role for WWOX loss of function in B cell neoplastic progression towards more aggressive phenotypes.

## Introduction

The *WWOX/FRA16D* chromosomal fragile locus is a hotspot for genomic instability (1-3). Consequently, *WWOX* is one of the most frequently deleted genes in cancer (4-7) and its expression loss is indicative of poor prognosis (2). Previous research has demonstrated increase incidence of B cell lymphomas in hypomorphic *Wwo*x mice (8) and in mice heterozygous for a functional *Wwox* allele upon carcinogen exposure (9), collectively indicating an increased vulnerability of B cells to neoplastic transformation upon *Wwox* deficiency. In humans, genetic alterations and depletion of *WWOX* have been linked to the pathogenesis of B cell malignancies such as certain B cell lymphomas (10-14) and multiple myeloma (MM) (15). *WWOX* is frequently deactivated by copy number loss and exhibits structural variations in B cell lymphomas (10-12,14). In MM, *WWOX* is frequently affected by deletions and breakpoints leading to translocation events commonly associated with poor prognosis, e.g. t(14;16) (16-20). Importantly, *WWOX* has recently been identified among a small number of cancer genes displaying complete loss of function in MM cases from the MMRF CoMMpass study, further supporting a potential role for *WWOX* in MM (21). Additionally, promoter methylation affecting *WWOX* expression has been associated with pathogenesis and advancement of MM (22). All these findings underscore a potential relevant role for *WWOX* deficiency in the progression and prognosis of B and plasma cell cancers.

To understand the impact of WWOX loss of function in B cells, we previously developed a *Cd19 Wwox KO* mouse model, where *Wwox* was conditionally deleted early in B cell development. These mice exhibited significantly reduced survival and developed B cell tumors including lymphomas, plasma cell neoplasias, and monoclonal gammopathies (23). *Wwox* ablation in B cells also caused spontaneous translocations during class switch recombination and a shift from classical non-homologous end-joining (NHEJ) to the microhomology-mediated alternative-NHEJ pathway, which is associated with chromosome translocations and genomic instability (23). This work suggested a propensity for oncogenic transformation in B cells upon *Wwox* deficiency.

In this study, we aimed to further elucidate the role of *WWOX* in B cell neoplastic progression by crossing *Cd19 Wwox KO* mice with *Vk*MYC* transgenic mice, a known model of MYC driven plasma cell neoplasia (24). Deletion of *Wwox* in B cells from *Vk*MYC* mice (*Vk*MYC:Wwox KO*) significantly reduced survival, primarily due to promoting the development of extramedullary plasmablastic plasmacytomas (PBPs), plasmablastic lymphomas (PBLs) and B cell lymphomas (BCLs). These tumors displayed a high mutation burden, additionally, transcriptome profiling of bone marrow (BM) derived Cd138+ plasma cells and plasmablastic tumors (PBTs) revealed an enrichment of biofunctions associated with tumorigenesis and inflammation activation as consequence of *Wwox* deletion. Together, our findings demonstrate that B-cell-specific *Wwox* deletion in a myeloma mouse model leads to the development of aggressive plasmablastic tumors characterized by hypermutation, overexpression of Aid/Apobec family members and pro-inflammatory signatures, further supporting the role of *Wwox* loss of function as a relevant event in malignant progression towards more aggressive phenotypes.

## Materials and Methods

### Animals

All animal research was conducted in facilities accredited by the Association for Assessment and Accreditation of Laboratory Animal Care International at the University of Texas MD Anderson Cancer Center, and all research was approved by the Institutional Animal Care and Use Committee. Mice were bred and kept in a clean, modified-barrier animal facility, fed regular commercial mouse diet (Harlan Lab., Indianapolis, IN) under controlled light (12 L:12D) and temperature (20–24 °C). The procedure to develop *Vk*MYC* mice with AID-dependent *Myc* activation in germinal center B cells was provided elsewhere (24), here we used a derivative strain described as *Vk*MYCΔloxP* in which LoxP site flanking flanking the transgenic 3’ kappa enhancer have been removed thus retaining expression in the presence of Cre recombination as recently described (25). The protocol to generate *Wwox^loxP/loxP^* and *Cd19:Wwox KO* mice has been previously described (23,26). We crossed *Cd19:Wwox WT* and *Cd19:Wwox KO* mice (C57Bl/6 background) with *Vk*MYC* (C57BL/6 background) to develop *Vk*MYC:Wwox WT, Vk*MYC:Wwox HET,* and *Vk*MYC:Wwox KO* mice. Mice of both genders were used in all experimental procedures. Genotypes were confirmed by PCR using primers previously described (23,24).

### Histology and immunohistochemistry

Full necropsy was performed on all mice and samples from major organs and tumors (whenever available). Tissues were processed by formalin fixation, paraffin embedding, and hematoxylin and eosin (H&E) staining. Tumor samples were also analyzed by immunohistochemistry (IHC) following standard procedures and staining with anti-CD138 (Biolegend 142502) and anti-CD19 (CST Rabbit mAb # 90176) antibodies. Histological analyses were performed without prior knowledge of genotypes. Based on IHC staining pattern and cell morphology, tumors were classified following guidelines of the Bethesda classification of lymphoid neoplasms in mice and a more recent classification of mouse plasmacytomas (27,28).

### Serum Protein Electrophoresis (SPEP)

Blood samples were collected from the tail vein at various intervals throughout the lifespan of the mice, or by cardiac puncture in moribund mice and those that survived until the end of the experimental period. The samples were allowed to coagulate at room temperature, then centrifuged at 3000 x g for 10 minutes. A volume of 0.5 μL of sera was loaded onto precast QuickGels (Helena Laboratories, 3505T) and run on a QuickGel Chamber (Helena Laboratories, 1284) according to the manufacturer’s instructions.

### Purification of CD138+ plasma cells and CD19+ B cells from mouse bone marrow

CD138+ plasma cells were purified from *Vk*MYC:Wwox WT* (n=5), *Vk*MYC:Wwox HET* (n=3) and *Vk*MYC:Wwox KO* (n=5) mice BM using EasySep^TM^ mouse CD138 positive selection kit (18957, STEMCELL Technologies) following manufacturer’s protocol. Briefly, BM was collected from mouse tibia and femur bones, washed, and resuspended in cold PBS with 2% FBS and 1 mM EDTA. Based on the number of cells obtained, BM cells were incubated with an FcR blocker and antibody selection cocktail, followed by the addition of magnetic RapidSpheres^TM^. Samples were placed in a magnetic block to purify CD138+ BM cells and washed with cold PBS to remove impurities. The supernatant containing the remaining BM cells was used to purify CD19+ B cells using EasySep^TM^ mouse CD19 positive selection kit II (18954, STEMCELL Technologies) following the manufacturer’s protocol, to be used as a control or source of B cells in other experimental procedures.

### RNA sequencing and data analysis

RNA-Seq and data analysis was done as previously described (29,30). Briefly, total RNA was extracted from *Vk*MYC:Wwox WT*-BM (n=3), *Vk*MYC:Wwox KO*-BM (n=3), and *Vk*MYC:Wwox KO*-PBT (n=4) samples using TRIzol reagent (Invitrogen) following the manufacturer’s protocol. RNA concentration and integrity were measured on an Agilent 2100 Bioanalyzer (Agilent Technologies), and samples with RNA integrity number over 8.0 were used for sequencing. Directional mRNA-seq libraries were constructed using the ScriptSeq v2 RNA-Seq Library Preparation Kit (Epicentre). We performed 76 nt paired-end sequencing on an Illumina HiSeq3000 platform, obtaining ∼40 million tags per sample. Short reads were mapped to the mouse reference genome (mm10) using TopHat v2.0.10. We employed several R/Bioconductor packages to calculate gene expression abundance at the whole-genome level using the aligned records (BAM files). Differentially expressed genes between *Vk*MYC:Wwox WT*-BM vs *Vk*MYC:Wwox KO*-BM and *Vk*MYC:Wwox KO*-BM vs *Vk*MYC:Wwox KO-*PBT were identified using DESeq2 based on normalized log2 count per million values (31). Data integration and visualization of differentially expressed transcripts (FDR < 0.05) were done with Multiexperiment Viewer Software. Functional enrichment analysis of dysregulated transcripts was performed using Ingenuity Pathway Analysis (IPA, QIAGEN Inc.)

### Whole exome sequencing and data analysis

DNA from *Vk*MYC:Wwox WT*-BM (n=5), *Vk*MYC:Wwox HET* (n=3), *Vk*MYC:Wwox KO*-BM (n=5), *Vk*MYC:Wwox KO*-PBT (n=4), and one normal C57Bl6/J mice liver samples were purified using DNeasy Blood and Tissue Kit (Qiagen). High-quality genomic DNA (250 ng) with 260/280 ratios greater than 1.8 were processed for library preparation using the Twist Exome Library Preparation Kit (Twist Bioscience) following the manufacturer’s instructions. Exome capture was performed using the Twist Exome Capture Kit (Twist Bioscience) and 100nt paired-end sequencing was done using Illumina NovaSeq 6000 with an estimated 200X coverage per sample. Image analysis, base-calling, and error calibration were performed using Illumina genome analysis pipeline at an average sequencing depth of 464 per sample. Sequenced 100nt paired-end reads were aligned against the mouse reference genome (mm10) using BWA v0.7.17 and marked for duplicates using Picard v2.27.4 (https://github.com/broadinstitute/picard/releases). Base quality recalibration was carried out using the GATK suite, and MuTect2 (GATK v4.4) was employed to identify single-nucleotide variants (SNVs) and insertions/deletions (INDELs). Following previously described methods (32), variants were filtered based on allele frequencies and compared against the Mouse Genome Project database of known germline variants (33). The filtered variants were annotated using the Variant Effect Predictor (release 96) (34), and further filtered by functional consequence. Chromosomal regions with copy number alterations were identified using CNVkit (version 0.9.9) (35). Mutational signatures and tumor mutational burden were computed using the SigProfiler Assignment resource (36) according COSMIC signature database (https://cancer.sanger.ac.uk/signatures/) based on non-synonyms somatic variants. Data integration and visualization of the mutational signature contribution across tumors were done with the MultiExperiment Viewer software (MeV v4.9).

### Quantitative RT-PCR

Quantitative RT-PCR was performed using total RNA extracted from purified Cd19+ B cells as previously described (30). Briefly, cDNA was synthesized using High-Capacity cDNA Reverse Transcription Kit (Applied Biosystems) following manufacturer’s instructions. The relative expression level for specific genes was determined in triplicate by qRT-PCR using the SYBR Green-based method. After normalization to Gapdh RNA expression, the average fold change was calculated using the 2^-(ΔΔCt) method described elsewhere (37). Following primers were used for qRT-PCR-*Wwox* F-5′-CGAGAACGGACAAGTGTTTTT-3′, R-5′-CGGATTATCGTCCACGGTAAAT-3′ and Gapdh 5′-TGGCCTTCCGTGTTCCTAC-3′, R-5′-GAGTTGCTGTTGAAGTCGCA-3′.

### Statistical analysis for mouse survival and tumor incidence

Overall survival and M-spike incidence analyses were performed using SPSS statistical software. Log rank test was applied to determine statistical significance. Chi square test was performed to compare tumor incidence and Student’s t-test was performed to compare gene expression using GrapPad Prism V 10, p-values < 0.05 were considered significant.

### Data availability

The RNA sequencing data discussed in this study have been deposited in NCBI’s Gene Expression Omnibus (38) and are accessible through GEO Series accession number GSE288262. The whole exome sequencing data discussed in this study are available upon request from the corresponding author.

## Results

### *Wwox* ablation in B cells promotes plasmablastic tumor development in *Vk*MYC* mice

To investigate the role of *WWOX* in B cell neoplastic progression, we generated a mouse model with targeted *Wwox* deletion early in B cell development and *MYC* activation by crossing *Cd19: Wwox KO* (23) and a derivative strain of *Vk*MYC* mice (24,25). We validated effective deletion of *Wwox* in Cd19+ B cells purified from bone marrow of *Vk*MYC:Wwox WT* and *Vk*MYC:Wwox KO* mice by qRT-PCR. *Wwox* mRNA expression was negligible in Cd19+ B cells from *Wwox* KO mice (p<0.05), confirming successful deletion (Supplementary Figure S1a). The survival of *Vk*MYC:Wwox KO* mice was compared with control littermates wild-type for *Wwox* expression. Overall survival was plotted using the Kaplan-Meier method and statistically analyzed by the Log-rank test. Although most *Vk*MYC:Wwox KO* mice displayed an indolent disease course, they exhibited a significantly shorter mean survival of 660 days vs. 800 days observed in *Vk*MYC:Wwox WT* mice (Log-rank p=0.002) (Figure 1).

**Figure 1.**
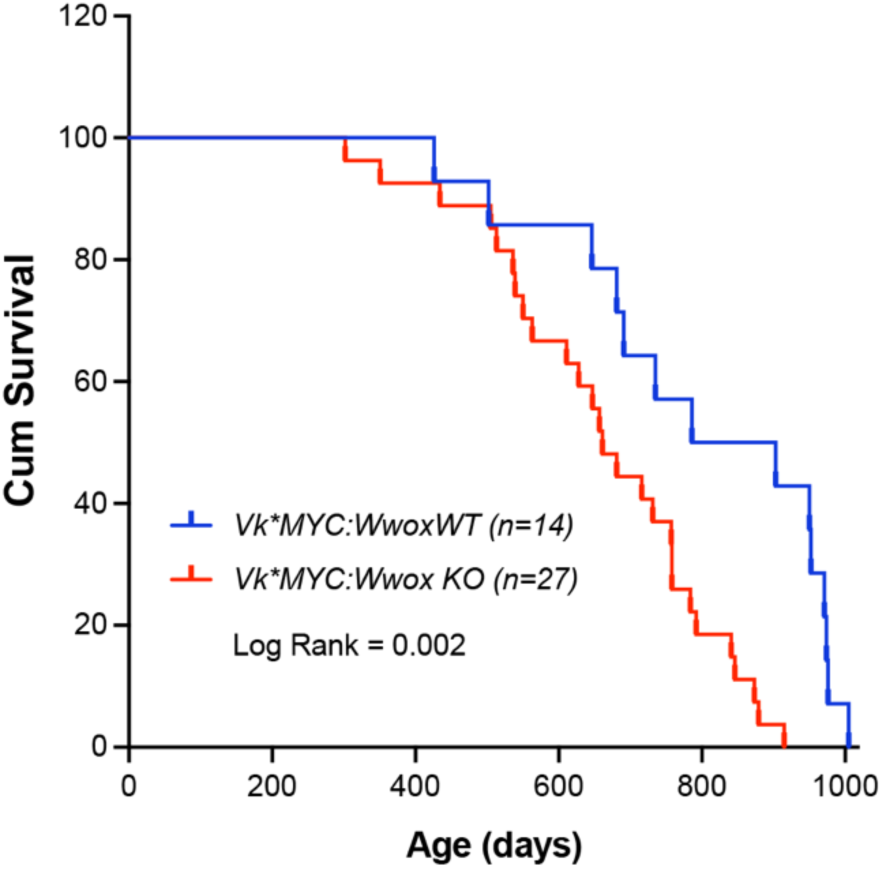
*Wwox* ablation in B cells reduces overall survival in *Vk*MYC* mice. Kaplan-Meier survival analysis comparing *Vk*MYC:Wwox KO* (red line, n=27) and *Vk*MYC:Wwox WT* (blue line, n=14) mice. *Vk*MYC:Wwox KO* mice showed significantly reduced survival compared to the WT group (Log-rank [Mantel-Cox] p-value = 0.002). The mean survival was 660 days for the *Wwox KO* group, compared to 800 days for the *Wwox WT* group.

We noted that 63% (17 out of 27) of *Vk*MYC:Wwox KO* mice developed intra-abdominal tumors with the characteristics of plasmablastic tumors and B cell lymphomas, particularly affecting mesenteric and peripancreatic lymph nodes (Figure 2a). This incidence was significantly higher than that observed in the *Vk*MYC:Wwox WT* group, where only 28.5% (4 out of 14) displayed intra-abdominal tumors (p = 0.036) (Figure 2a). Based on histopathology evaluation and IHC staining these tumors were broadly classified into, PBPs (n=9), PBLs (n=5) and BCLs (n=3) (Figure 2b, Supplementary Table S1). Interestingly, IHC revealed that histology samples from several *Vk*MYC:Wwox KO* extramedullary plasmacytomas (i.e., PBPs) displayed large areas characterized by the presence of Cd138+ plasma cells with staining to cell membrane but also to small single polarized foci (Figure 2c). This striking intracellular pattern of Cd138 staining has been previously described as resembling ‘uropods’ shown to facilitate myeloma cells aggregation and adhesion to the tumor microenvironment. Furthermore, myeloma cells with high proportion of cells with Cd138 staining uropods were shown to display high migratory potential, implying that they represent more aggressive tumor forms (39), which indeed agrees with the histology observed. Abundant number of aberrant mitotic figures, enlarged nuclei, and multiple prominent nucleoli were also characteristic of all PBPs (Figure 2c).

**Figure 2.**
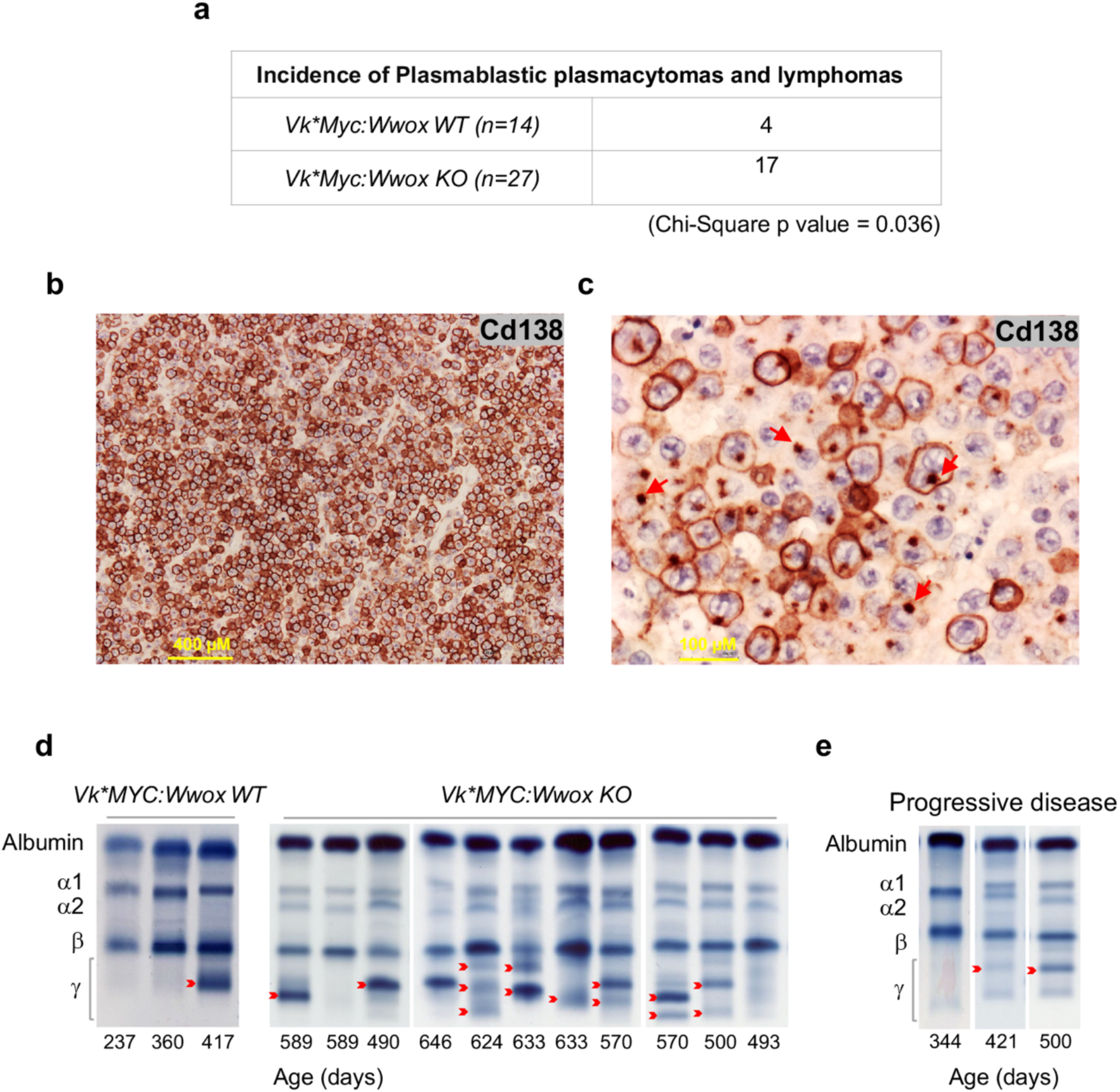
Increased tumor incidence in *Vk*MYC:Wwox KO* mice with high Cd138+ expression and uropod formation in plasmablastic plasmacytomas. **(a)** A significantly higher number of *Vk*MYC:Wwox KO* mice (17 out of 27) developed PBTs and BCLs compared to *Vk*MYC:Wwox WT* mice (4 out of 14), as determined by Chi-Square analysis (p-value = 0.0366). **(b)** Low-magnification, photomicrograph of extramedullary plasmablastic plasmacytoma histology section from a *Vk*MYC:Wwox KO* mouse immunostained with Anti-Cd138 antibody. Scale bar on lower left indicates 400 µM in length. **(c)** High-magnification photomicrograph of histology section from another *Vk*MYC:Wwox KO* PBP displaying areas of cells with immunostained uropod like structures, i.e., polarized intracellular Cd138 staining to small membrane protrusions (red arrows). Scale bar on lower left represents 100 µM in length. **(d) Representative SPEP of serum samples from *Vk*MYC:Wwox* WT and KO mice.** Red arrows indicate representative M-spikes. **(e)** Representative longitudinal follow up of serum samples analyzed by SPEP collected from the same mouse through time, displaying progressive disease with a marked increase in M-spike intensity over time.

A characteristic of monoclonal gammopathies such as multiple myeloma, plasmacytomas, and other plasma cell neoplasms is the secretion of immunoglobulin (Ig), which manifests as Ig monoclonal bands (M-spikes) in SPEP. We performed a longitudinal follow-up with SPEP in 24 out of 27 *Vk*MYC:Wwox KO* mice, as 3 mice died before the SPEP follow-up could be completed. All 24 *Wwox KO* mice (100%) showed detectable M-spikes, accompanied by a progressive increase in the intensity of secreted gamma globulin as mice aged, indicating disease progression (Figures 2d and e, Supplementary Figure S2). M-spikes were also observed in all *Vk*MYC:Wwox WT* mice, with 14 out of 14 (100%) showing secreted gamma globulin. Overall, there was no difference in the age of M-spikes detection when comparing *Vk*MYC:Wwox KO vs. WT* counterparts (Supplementary Figure S2).

### Pro-inflammatory and tumorigenic transcriptomic signatures in plasma cells with *Wwox* deletion

To gain insight into transcriptome changes associated with B cell specific *Wwox* deletion in *Vk*MYC* mice, we performed RNA-seq analysis of BM derived Cd138+ plasma cells from *Vk*MYC:Wwox KO* and *Vk*MYC:Wwox WT* mice. The differential gene expression (DGE) dataset identified a total of 87 dysregulated genes (FDR < 0.05) in *Vk*MYC:Wwox KO* vs. *Vk*MYC:Wwox WT* Cd138+ BM cells (52 upregulated and 35 downregulated genes) (Supplementary File S1). Unsupervised hierarchical clustering using DGE profiles robustly segregated *Wwox WT* from *Wwox KO* BM samples (Figure 3a). Among the genes showing differential regulation, a number of immune regulatory and pro-tumorigenic genes, such as *Foxq1*, *Vegfa*, *Jun*, and *Tgfb2*, were significantly upregulated in the *Wwox* KO group, as detailed in Supplementary File S1.

**Figure 3.**
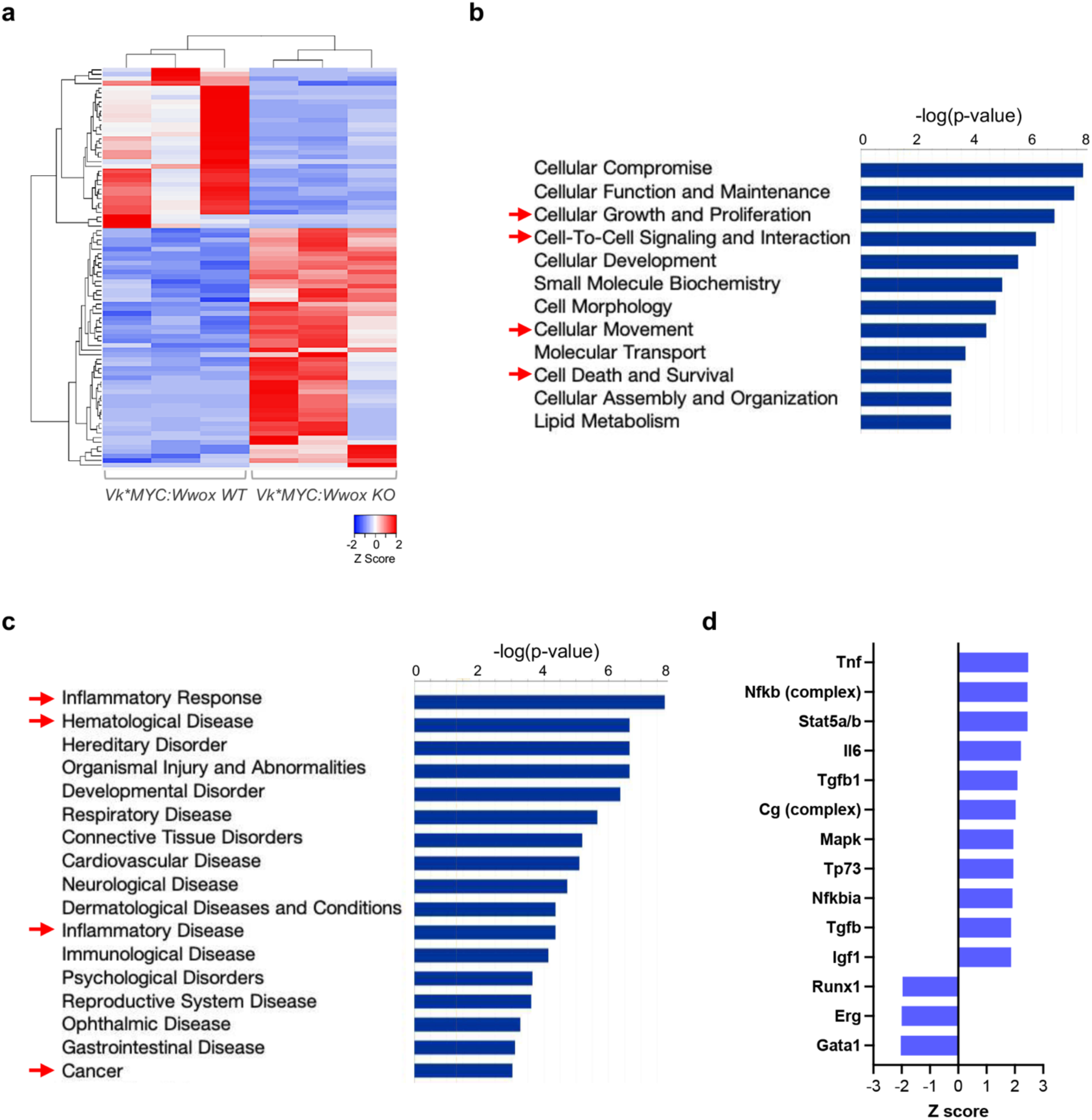
*Vk*MYC:Wwox KO* Cd138+ plasma cells display pro-inflammatory and tumorigenic transcriptomic signatures. **(a)** Unsupervised clustering and heatmap of 87 differentially expressed genes (FDR < 0.05) comparing *Vk*MYC:Wwox WT* and *Vk*MYC:Wwox KO* Cd138+ plasma cells (n=3 mice/group). Red or blue colors indicate differentially up or downregulated genes, respectively. Mean signals were background corrected and log2 transformed. **(b, c)** Bar graph depicting enrichment of biofunctions associated with tumorigenic phenotypes (**b**) and disease processes (**c**) (highlighted by red arrows), *driven by dysregulated gene expression in Vk*MYC:Wwox KO* plasma cells (Z-score cutoff ±2, p < 0.05). (**d**) Bar graph showing top-activated and inhibited upstream regulators as per IPA (Z-score cutoff ±2, p value < 0.05) based on DGE profiles in *Vk*MYC:Wwox KO compared to the control group*.

IPA-based functional annotation of differentially expressed genes in *Vk*MYC:Wwox KO* plasma cells revealed enrichment of biofunctions related to tumorigenic phenotype such as ‘Cellular growth and proliferation’, ‘Cellular movement’, ‘Cell death and survival’, and ‘Cell-to-cell signaling and interaction’ (Figure 3b). Additionally, enriched disease processes terms in the *Wwox KO* group included ‘Inflammatory response’, ‘Hematological disease’, ‘Inflammatory disease’, and ‘cancer’ (Figure 3c). Based on DGE profiles we also identified top-activated and inhibited upstream regulators (Z-score cutoff ±2, p value < 0.05). Tnf (activation Z-score 2.47) was identified as the topmost activated regulator along with other inflammatory pathway factors like the Nfkb complex and Il6. Furthermore, Tnf based regulation of several differentially expressed genes in our dataset pointed to ‘proliferation of tumor cells’ as a significant outcome effect in *Vk*MYC:Wwox KO* Cd138+ BM cells (Supplementary Figure S3). In addition, pro-tumorigenic signaling pathway molecules such as Mapk and Tgfb1, along with transcription factor Stat5a/b were among other significantly activated upstream regulators, while Runx1, Erg, and Gata1 were found as inhibited regulators (Figure 3d). These observations strongly link transcription factors, signaling pathways, and inflammatory changes that contribute to a tumorigenic phenotype with the specific deletion of *Wwox* in B cells from *Vk*MYC* mice.

### Plasmablastic tumors displayed highly enriched inflammatory gene expression signatures

We further compared the transcriptome profiles and investigated the differences between Cd138+ BM plasma cells and PBTs from *Vk*MYC:Wwox* KO mice. This analysis identified 2355 differentially regulated genes (FDR < 0.05) in *Vk*MYC:Wwox KO-*PBT vs *Vk*MYC:Wwox* KO*-* BM cells (1389 upregulated and 966 downregulated genes) (Supplementary File S1). Unsupervised hierarchical clustering of the DGE dataset showed clear segregation of BM plasma cells and plasmablastic tumors (Figure 4a). In *Wwox* KO plasmablastic tumors, a noteworthy observation was the pronounced upregulation of genes implicated in facilitating cancer cell migration, invasion, and metastasis such as *Ccl21a*, *Cilp*, *Mmp3*, and *Pdpn*, exhibiting more than a nine-fold difference in expression (Log2 FC range: 9.09 – 12.53) (Supplementary File S1). Interestingly, the *Aicda* gene (activation induced cytidine deaminase, also known as *Aid)*, which encodes a key enzyme expressed in germinal centers involved in somatic hypermutation, gene conversion and class switch recombination of Ig genes, was found to be overexpressed 11 Log2 fold higher on average in plasmablastic tumors vs. BM cells (Supplementary Figure S4). Likewise, *Utf1*, a gene linked with stem cell properties and cancer stemness, was significantly upregulated in *Wwox KO*-PBT samples (Log2 FC =9.32) (Supplementary File S1). Similar to observations in the *Wwox KO* vs *WT* BM comparison, transcriptome profiling of *Wwox KO-*PBT vs. *Wwox KO-* BM samples, also showed enrichment of ‘Cell death and survival’, ‘Cellular movement’, ‘Cellular growth and proliferation’, and ‘Cell-to-cell signaling and interaction’ tumorigenic biofunctions and ‘Cancer’, ‘Hematological disease’, ‘Inflammatory disease’, and ‘Inflammatory response’ as significantly enriched conditions (Figures 4b and c). However, the number of deregulated genes and magnitude of statistical significance for all enriched biofunctions and disease processes were far more pronounce in *Vk*MYC:Wwox KO-*PBT (p-value range 4.28E-40 – 1.69E-19) in comparison to *Vk*MYC:Wwox KO-*BM (p-value range 2.06E-07 – 1.19E-03) (Figures 4d and e). Interestingly, one of the significant regulator effects identified in *Wwox KO*-PBT is based on the upstream regulator Ccl2 (also known as Mcp10), a potent chemoattractant cytokine with an activation Z-score of 4.64 and capable of upregulating multiple target genes in the DGE dataset, ultimately leading to increased ‘Homing of cells’ and ‘Inflammatory response’ (Supplementary Figure S5). The enrichment of the same biofunctions and diseases suggests consistency between *Vk*MYC KO-*BM cells and PBTs. However, markedly higher number of differentially expressed genes and magnitude of statistical significance in the enrichment analysis strongly indicate a more pronounced tumorigenic and inflammatory phenotype in PBTs, likely driving the plasmablastic/anaplastic progression of malignant plasma cells.

**Figure 4.**
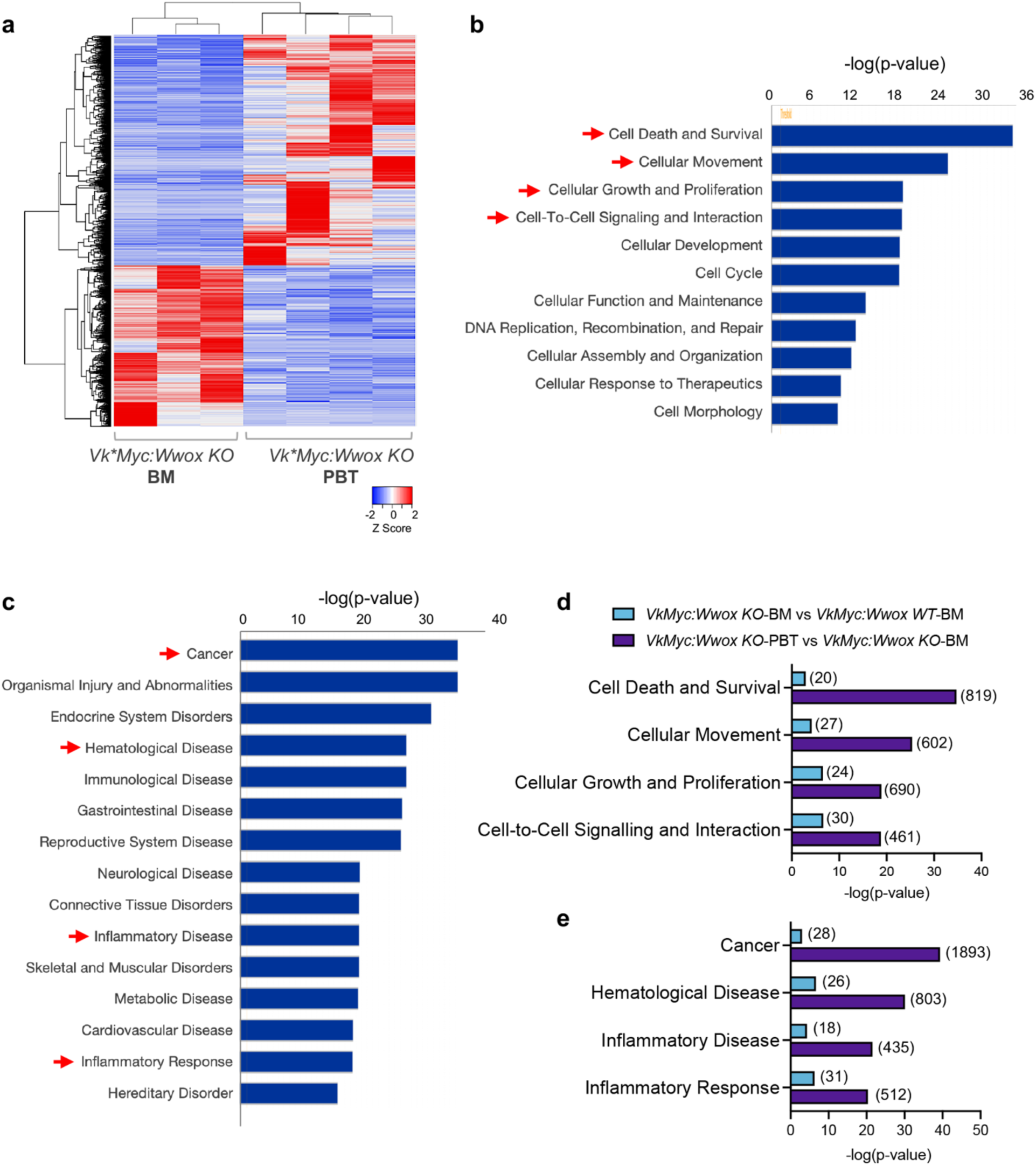
Highly enriched inflammatory gene expression signatures in *Vk*MYC:Wwox KO* plasmablastic tumors. (**a**) Unsupervised clustering and heatmap of 2355 differentially expressed genes (FDR < 0.05) comparing *Vk*MYC:Wwox KO*-BM (n=3) and *Vk*MYC:Wwox KO*-PBT (n=4) groups. Red or blue colors indicate differentially up or downregulated genes, respectively. Mean signals were background corrected and log2 transformed. **(b, c)** Bar graph depicting enrichment of biofunctions associated with tumorigenic phenotypes (**b**) and disease processes (**c**) (highlighted by red arrows), *driven by dysregulated gene expression in Vk*MYC:Wwox KO* plasmablastic tumors (Z-score cutoff ±2, p < 0.05). (**d**) Bar graph showing the number of deregulated genes and the magnitude of statistical significance for enriched biofunctions and disease processes in *Vk*MYC:Wwox KO-PBT vs Vk*MYC:Wwox KO-BM* and *Vk*MYC:Wwox KO-BM vs Vk*MYC:Wwox WT-BM* groups. The enrichment was more pronounced in *Vk*MYC Wwox KO-PBT vs Vk*MYC Wwox KO-BM (p-value range: 4.28E-40 to 1.69E-19) compared to Vk*MYC:Wwox KO-BM vs Vk*MYC:Wwox WT-BM* (p-value range: 2.06E-07 to 1.19E-03). Numbers in parentheses beside each bar indicate the number of differentially expressed genes in each category.

### *Vk*MYC:Wwox KO* plasmablastic tumors display hypermutation profiles and multiple cancer driver mutations

By means of whole exome sequencing we analyzed the mutational profile of Cd138+ BM cells from *Vk*MYC:Wwox KO*, *HET*, and *WT* mice, and from *Vk*MYC:Wwox KO* PBT tissue samples. Among the four PBT samples analyzed, three were classified as PBPs, while one was identified as a PBL. Non-synonymous single nucleotide variants (SNV) and INDELs were analyzed, and variant effect predictor and SIFT impact scores were employed to identify relevant protein mutations affecting cancer driver genes across all sample groups. No cancer driver mutations were detected in any of the *Vk*MYC:Wwox WT-*BM samples, while two of the *Wwox HET*-BM and one *Wwox KO-*BM displayed deleterious mutations affecting some cancer genes (Figure 5). The scarcity of cancer driver mutation in most of the BM samples regardless of genotype, suggests that the main driver is just *MYC* overexpression and is in agreement with recent observations (25). In contrast, all *Vk*MYC:Wwox KO* plasmablastic tumor samples exhibited multiple mutations affecting various cancer genes commonly associated with both multiple myeloma and B-cell lymphomas. Notably, the highest number of mutations was observed in the *Wwox KO* PBL sample, which harbored 21 distinct cancer driver mutations.

**Figure 5.**
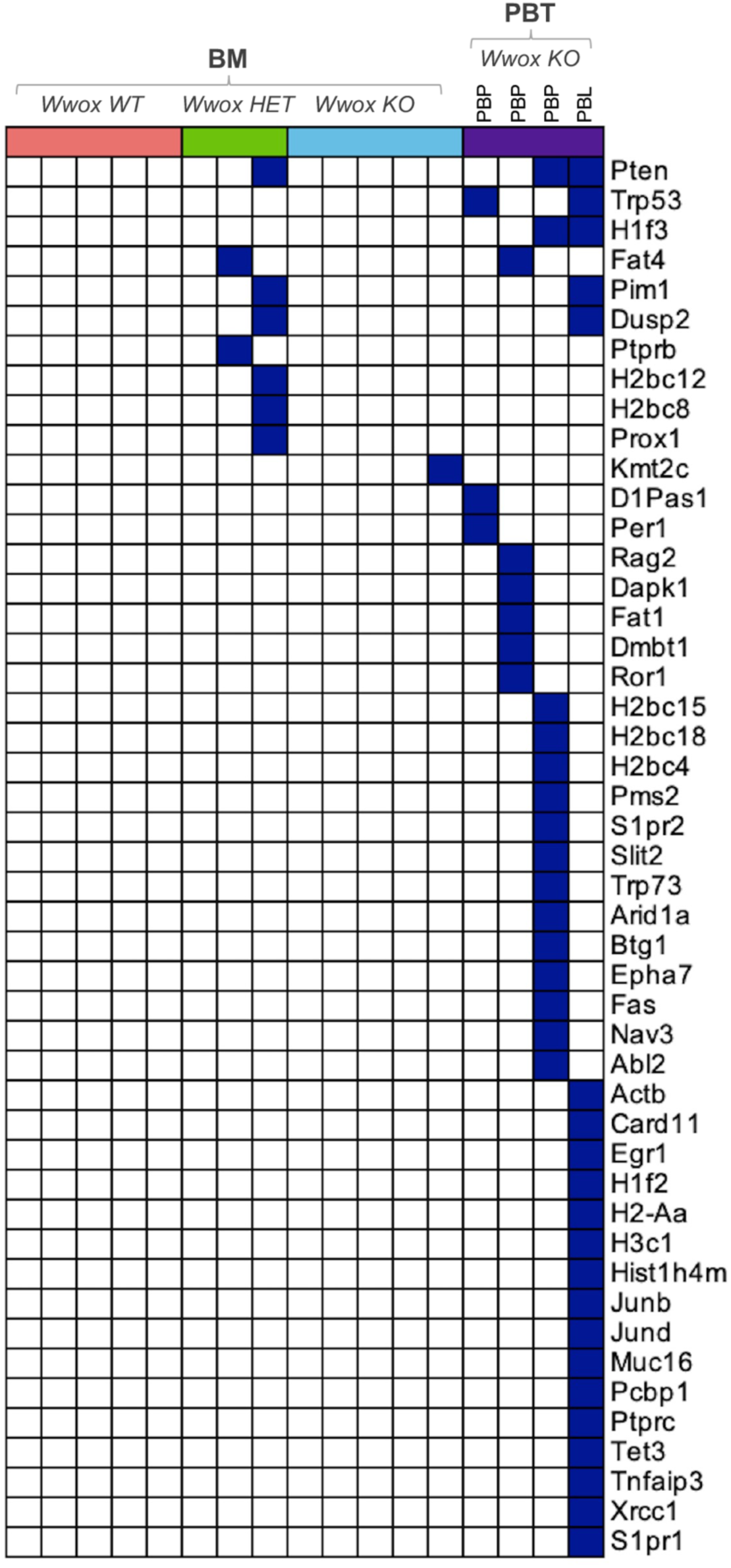
Somatic, non-synonymous SNVs in cancer driver genes in *Vk*MYC:Wwox KO* plasmablastic tumors. Oncoplot displaying non-synonymous SNVs in cancer driver genes identified through exome sequencing of Cd138+ BM plasma cells and *Vk*MYC:Wwox KO PBTs. Each column represents an individual mouse, grouped as indicated at the top of the plot, and each row corresponds to mutations detected in a specific gene for these mice. In the Vk*MYC:Wwox KO*-PBT group, the first three columns represent PBPs, while the last column represents a PBL. Blue squares indicate the presence of mutations in the corresponding gene for each sample.

Among all the identified mutations, the tumor suppressor *Pten* emerged as the most frequently mutated gene, detected in one *Wwox HET*-BM sample and in two *Wwox KO* plasmablastic tumors— one in PBP and the other in the PBL. Other mutated genes, common to both BM and PBT groups, included the tumor suppressor *Fat4*, identified in one *Wwox HET*-BM and one *Wwox KO*-PBP sample. Additionally, mutations in the oncogene *Pim1* and the cell signaling gene *Dusp2* were both found in one *Wwox HET*-BM sample and the *Wwox KO*-PBL. Notably, all these genes have been previously reported as multiple myeloma drivers (40-42) and within the *Vk*MYC* model (25) (Figure 5). Among other recurrent mutations in *VkMyc:Wwox KO-*PBTs were the tumor suppressor gene *Trp53* and the histone gene *H1f3*, each present in 2 out of 4 samples, including both PBP and PBL. Several other histone genes, *H2bc12, H2bc8, H2bc15, H2bc18, H2bc4, H1f2, H2-Aa, H3c1, Hist1h4m*, were also found mutated, underscoring a potential disruption of chromatin dynamics and gene expression in these tumors. In addition, we identified mutations in other tumor suppressors including *Fat1, Dapk1,* and the *Ror1* oncogene. Both PBPs and PBL also showed mutations in DNA damage repair genes (*Arid1a, Pms2, Xrcc1*) and genes involved in proliferation and cell cycle regulation (*Junb, Jund, Per1, Egr1, Btg1, Tet3*), further highlighting genomic instability. Moreover, several mutations were detected in cell signaling genes, suggesting broad perturbations across oncogenic networks (Figure 5). Mutations in the described cancer drivers have been reported not only in MM (40-42) but also in B-cell lymphomas as well (43).

We also performed single based substitution (SBS) signature analyses to characterize the most represented signatures and uncover underlying mutagenic processes. All samples exhibited signatures SBS1, associated with spontaneous 5-methylcytosine deamination, and SBS5, known for its clock-like mutation profile (Figure 6a, Supplementary Figure S6). Both signatures are associated with age-related mutational processes, with SBS5 further linked to DNA replication stress. The consistent presence of these signatures indicates a background of aging-related mutations and ongoing replication stress across all samples. The most significant finding from our analysis was the detection of DNA repair related mutational activity, marked by the presence of SBS26 and SBS30 (Figure 6a, Supplementary Figure S6). SBS26, associated with defective DNA mismatch repair, was observed in samples from all BM groups including two samples from the HET and one from the KO groups (Figure 6a, Supplementary Figure S6). Additionally, SBS30 linked to defective base excision repair (BER), was observed only in *Vk*MYC:Wwox KO*-PBPs (2 out of 3 samples) (Figure 6a, Supplementary Figure S6). Additionally, we identified SBS32 and SBS87, both associated with thiopurine family drugs exposure. These drugs are known to be mutagenic via the integration of the final metabolite 6-thioguanine into DNA. Importantly, one route of escape of thiopurines cytotoxicity is by inactivation of mismatch repair mechanisms (44). Perhaps and like the described SBS26 and SBS30 associated with defective BER and mismatch repair, the finding of SBS32 and SBS87 signatures also reflects associations with DNA mismatch repair mechanisms. Additionally, we detected the SBS85 signature, which reflects the indirect effects of Aid activity in lymphoid cells. Other signatures, including SBS12, SBS37, and SBS40a, of unknown origin, were also detected (Figure 6a).

**Figure 6.**
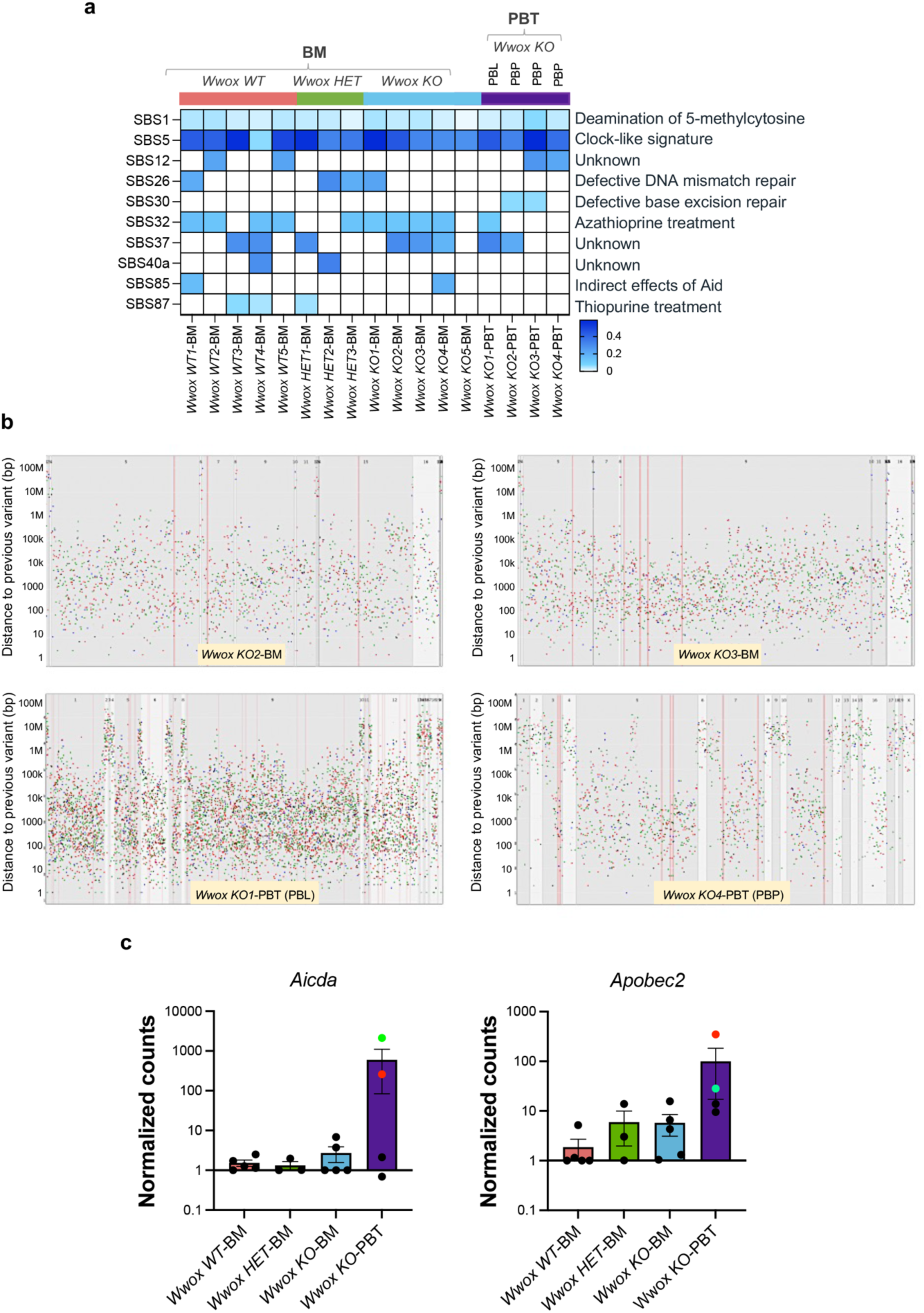
Distinct SBS signatures and the occurrence of kataegis-like clustered hypermutation profiles in *Vk*MYC:Wwox KO* BM and plasmablastic tumors. (**a**) Heatmap displaying the most frequent SBS mutational signatures identified in BM plasma cells and *Vk*MYC:Wwox KO* PBPs. The Y-axis represents the distinct SBS signatures detected in these samples, while the X-axis denotes the individual samples. *Each column represents an individual mouse, grouped as indicated at the top of the heatmap. In the Vk*MYC:Wwox KO*-PBT group, the first column represent PBL, while the remaining three columns represents a PBPs. Color intensity on the heatmap indicates the strength of association with each SBS signature. (**b**) Rainfall plots of all the mutations across the genome of representative samples from *Vk*MYC:Wwox KO* BM and PBTs demonstrating a widespread distribution of clustered hypermutations resembling kataegis. The Y-axis represents the distance of each mutation from the preceding mutation. Different colors represent the different types of the mutation substitutions. Clustering of mutations closer to the X-axis indicates a smaller distance between mutations, reflecting hypermutation. (**c**) Bar graphs illustrating the average normalized counts of *Aicda* and *Apobec2* mRNA expression in *Vk*MYC:Wwox WT* (n=5)*, HET* (n=3), *KO* (n=5) BM and *Vk*MYC:Wwox KO* PBT (n=4) samples. Each data point represents the expression level for an individual mouse, with error bars indicating the mean ± SEM. The green data point indicates *Aicda* and *Apobec2* expression in the *Vk*MYC:Wwox KO1*-PBT (PBP), while the red data point indicates expression in the *Vk*MYC:Wwox KO4*-PBT (PBL).

Interestingly, 2 out of 5 *Vk*MYC:Wwox KO*-BM samples and 2 out of 4 *Vk*MYC:Wwox KO*-PBTs, including one PBP and one PBL, displayed a widespread distribution of clustered mutations resembling kataegis, with a broader mutational pattern consistent with hypermutated genomic regions (Figure 6b, Supplementary Figure S7). This profile was particularly prominent in the PBL, which exhibited an exceptionally pronounced hypermutated pattern (Figure 6b). Mutation clusters and hypermutation in cancer have been hypothesized to result from the activity of AID/APOBEC editing deaminases (44). Consistent with this hypothesis, expression analysis of *Aid/Apobec* family members in *Vk*MYC:Wwox KO*-PBT samples revealed marked overexpression of *Aicda* and *Apobec2* in the PBP and PBL tumors exhibiting hypermutation (Figure 6c), while no notable differences were found in *Apobec1* and *Apobec3* expression (Supplementary Figure S8). Overall, these findings and those of mutational signatures described above, suggest that *Wwox* deletion impacts DNA repair processes with *Wwox KO* samples exhibiting mutational profiles compatible with DNA repair deficiencies and hypermutation.

### Extensive chromosomal copy number alterations are characteristic of *Vk*MYC:Wwox KO* plasmablastic tumors

We next analyzed CNVs by comparing the genomic profiles of BM and PBT samples. Similar to SNV analyses, copy number alterations were relatively low and comparable between all BM sample groups again pointing to a rather indolent disease status mostly driven by *MYC* overexpression (Figure 7a, Supplementary Figures S9a, b, and c, Supplementary Table S2). In contrast, both copy number gains and losses were significantly elevated in *Vk*MYC:Wwox* KO plasmablastic tumors (Figures 7a and b, Supplementary Table S2).

**Figure 7.**
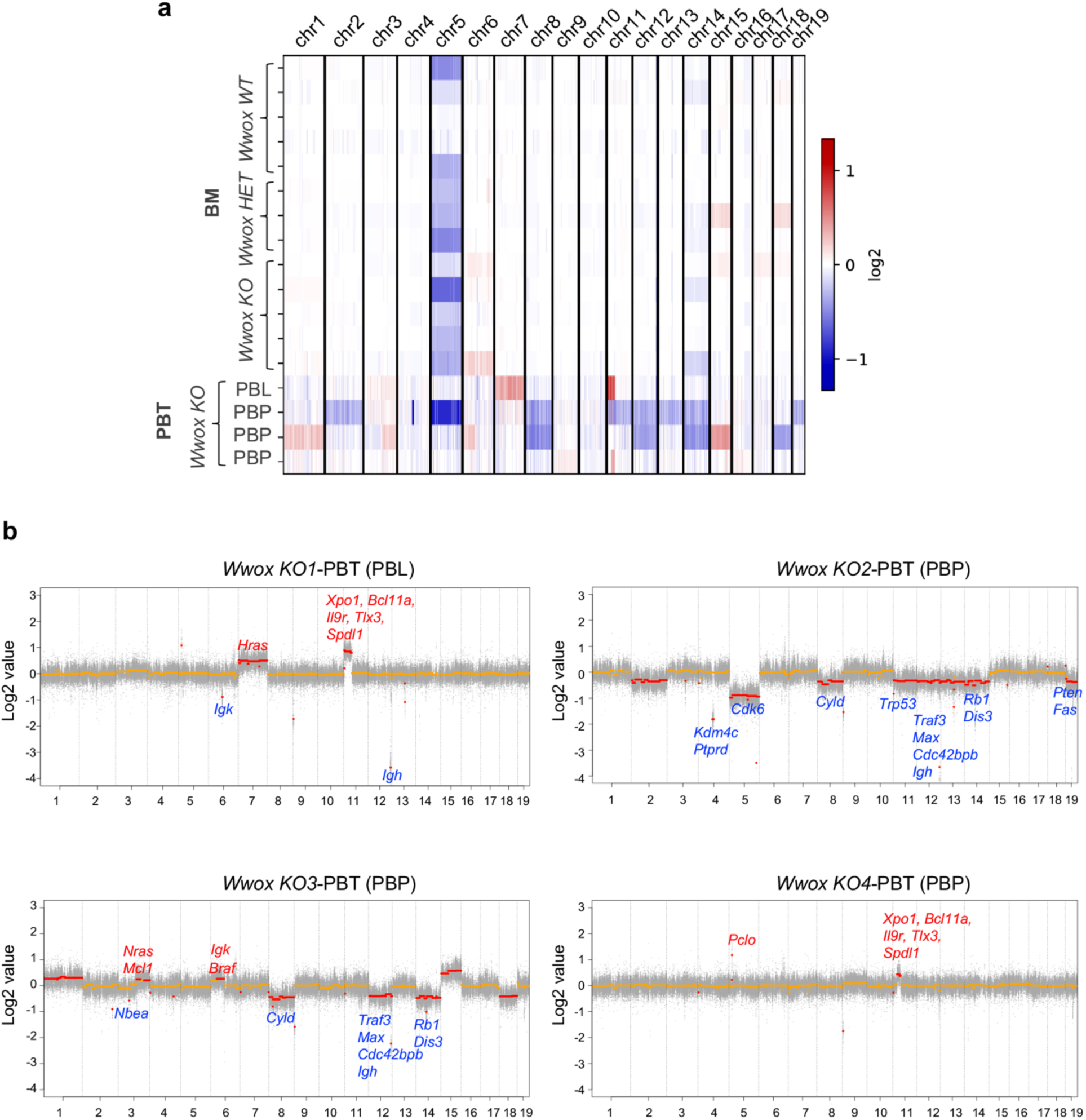
*Vk*MYC:Wwox KO* PBTs display recurrent focal and large copy number alterations. (**a**) Heatmap illustrating copy number abnormalities in BM plasma cells and *Vk*MYC:Wwox KO* PBPs. The Y-axis represents individual samples, while the X-axis displays individual chromosomes, separated by solid lines. Color coding indicates copy number losses or gains at various chromosomal loci, with blue representing copy number loss and red indicating copy number gain. The intensity of the colors reflects the magnitude of the copy number alterations. (**b**) Scatter plots from *Vk*MYC:Wwox KO*-PBT samples illustrate widespread and pronounced copy number alterations across all samples. Focal and large copy number alterations are represented by red markers; those above zero indicate copy number gains, while markers below zero signify copy number losses. The names of the genes affected by specific copy number alterations are displayed near the corresponding alterations, with red indicating copy number gains and blue indicating copy number losses for the respective genes.

Monosomy or large deletions of chr.5 were observed across all groups, with 2 out of 5 *Vk*MYC:Wwox WT*-BM, 3 out of 3 *Vk*MYC:Wwox HET*-BM, and 3 out of 5 *Vk*MYC:Wwox KO*-BM (Supplementary Figures S9a, b, and c). Additionally, *Vk*MYC:Wwox KO* PBT case exhibited losses of chr.5, while two cases displayed small focal gains (Figure7b). Notably, such extensive chr.5 deletions have previously been implicated as drivers of tumor progression in the *Vk*MYC* myeloma model (25,45,46).

Focal deep deletions affecting chr.12 were identified in one sample from each BM group, while these CNVs were more frequent in PBTs, occurring in 3 out of 4 samples (two PBPs and one PBL), with two PBPs showing extensive chr.12 deletions (Figure 7b, Supplementary Figures S9a, b, and c). Focal deletions affecting chr.6 were also observed in 1 out of 5 *Wwox KO*-BM samples and the PBL out of 4 *Wwox KO*-PBTs. The described deletions primarily affected the *Igh* locus on chr.12 and the *Igk* locus on chr.6, both crucial for antibody diversity (Supplementary Table S2). Interestingly, the described chr.12 large deletions observed in 2 out 4 PBPs also impacted *Traf3*, *Max*, and *Cdc42bpb* genes (Figure 7b), all described among the handful of targets in addition to *Wwox* displaying ‘complete loss of function’ in the latest CoMMpass study (21). *Vk*MYC:Wwox KO* plasmacytomas also displayed large chr.14 losses in 2 out of 4 samples impacting two critical genes, *Dis3* and the tumor suppressor *Rb1*. (Figure 7b). Mouse chr.14 shows significant synteny with human chr.13q, known for being among the most common large chromosomal losses found in MM and other human lymphoid malignancies (45,47) and involving *DIS3* and *RB1* genes, both recognized as key drivers in MM (21). Large deletions affecting chr.8, which includes the *Cyld* locus, were also observed in 2 out of 4 *Vk*MYC:Wwox KO* PBPs (Figure 7b).

Additionally, an amplicon was detected affecting chr.11 in *Vk*MYC:Wwox KO*1 PBP and PBL samples involving the *Rel* oncogenic Nf-kb subunit, *Xpo1*, *Bcl11a*, *Il9r*, *Tlx3*, and *Spdl1* all of which play critical roles in lymphoid development and oncogenesis (48-52) (Figure 7b). A focal amplification affecting chr.6 and including the *Braf* locus was identified in one sample each from the *Vk*MYC:Wwox KO* BM and PBP groups, gene also displaying gain of function in MM (21) (Figure 7b, Supplementary Figure 9c). Among other significant alterations found in individual tumors was a large copy number loss on chr.19, found in one *Vk*MYC:Wwox KO*1 PBP, which contains the critical tumor suppressors *Pten* and *Fas*, both vital for regulating cell survival and apoptosis (Figure 7b). Additional CNVs identified in individual PBTs were focal losses in *Kdm4c*, *Ptprd*, *Trp53*, as well as gains of chromosomes containing oncogenes such as *Hras*, *Nras*, *Mcl1*, and *Pclo* oncogenes (Figure 7b).

## Discussion

We generated a *Vk*MYC:Wwox KO* mouse cancer model, harboring two oncogenic alterations commonly found in B and plasma cell cancers: *Myc* activation and *Wwox* deletion. The *Vk*MYC:Wwox KO* mice exhibited significantly reduced survival rates when compared with *Vk*MYC:Wwox WT* counterparts, demonstrating that B cell-specific Wwox deficiency in mice with *Myc* activation leads to the development of more lethal cancer phenotypes. These mice displayed a significant high incidence of plasmablastic tumors characterized by distinct histopathological features and immunohistochemical staining patterns compatible mostly with aggressive extramedullary plasmablastic plasmacytomas and plasmablastic lymphomas. A particularly interesting finding in plasmacytomas was the presence of clustered Cd138 staining to polarized cellular foci previously described as resembling uropods in *Vk*MYC:Wwox KO* plasmacytoma cells. This phenomenon is known to correlate with heightened migratory potential and aggressiveness in myeloma cells and thus highlights the potential role of Wwox deletion in fostering a more aggressive tumor phenotype (39).

Among the most important observations from these studies was the significant activation of pro-inflammatory pathways in *Wwox KO* BM cells and plasmablastic tumors. Transcriptome profiling demonstrated significant enrichment of inflammatory signatures, with key regulators such as Tnf, Nf-kb, Il6, and Ccl2 prominently involved, highlighting the central role of inflammation in driving tumor progression in Wwox deficient models. These findings are in agreement with previous observations across cancers, CNS disorders, and various inflammatory conditions indicating a significant role for WWOX in inflammation regulation due to direct associations with the NF-kB and IL-6/JAK2/STAT3 signaling pathways (29,30,53-60). Chronic inflammation is well known to be crucial in cancer development, fostering an immunosuppressive environment, promoting cell proliferation, survival, migration and contributing to genomic instability through cytokines and chemokines (61) (62). NF-kB is a known master regulator of inflammatory response transcriptionally controlling expression of proinflammatory cytokines (e.g. IL1, IL6, IL8, TNF) as well as anti-apoptotic genes (e.g. BCL2, BCLXL) among others (63). Furthermore, production of cytokines like TNF and IL6 and growth factors that promote tumor progression, have been associated with more aggressive forms of B cell lymphomas and multiple myeloma (62).

The exome-seq analysis identified a single oncogenic mutation and minimal copy number alterations in *Vk*MYC:Wwox KO*-BM cells, indicating an indolent disease state in the bone marrow regardless of genotype and suggesting that *MYC* overexpression functions as the main oncogenic driver accompanied only by the common copy number losses affecting chr.5. In contrast, *Vk*MYC:Wwox KO* plasmablastic tumors exhibited multiple mutations and significant number of CNVs affecting key cancer driver genes, indicating increased genomic instability and a clear shift towards a more aggressive disease state. Mutation analysis identified various classical myeloma cancer target genes, including *Pten, Trp53, Pim1, Dusp2, Btg1, Egr1*, as well as various histone targets such as *H1f3, H1f2* and others that align with those recently reported in *Vk*MYC* derived models (25), thereby underscoring their relevance in hematologic malignancies, including B-cell lymphomas and multiple myeloma (64-66). Likewise, CNV findings aligned with the recent comprehensive MMRF CoMMpass study of newly diagnosed MM patients that led to the identification of only 12 target genes with recurrent and complete loss of function in >2% of the cohort that include and coincide with target losses in our study such as *TRAF3, DIS3, RB1, CYLD, TP53, CDC42BPB* and indeed *WWOX* by study design in our case (21). Furthermore, the authors of this study described that although loss of one *WWOX* allele is expected in t(14;16) they frequently detected complete loss of function at approximately the same frequency of total loss of classical targets such as RB and TP53, indeed supporting a possible role for WWOX in MM (21). Similarly, copy number gains including *Braf* and *Ras* family oncogenes, identified in our PBPs and also described in the CoMMpass study were also found.

*Wwox KO* plasmablastic tumors also displayed mutations of various DDR genes, mutational signatures associated with defective DNA damage repair and overexpression of Aid/Apobec gene family members, known for inducing somatic hypermutations through cytosine deamination all processes contributing to genomic instability, tumorigenesis and described in MM (67,68). Notably, we and others have shown that loss of WWOX function significantly impairs DNA repair efficiency, resulting in the accumulation of DNA damage that triggers genomic instability, a hallmark of cancer progression (3,23,69-71).

In conclusion, the described findings strongly support that the loss of WWOX function is a key driver for the development of aggressive plasmablastic phenotypes, primarily through two mechanisms: escalating genomic instability and fostering a pro-tumorigenic inflammatory microenvironment.

## Acknowledgement

Authors would like to thank the University of Texas MD Anderson Cancer Center (UTMDACC) Research Animal Support Facility and Laboratory Animal Genetic Services for animal support and Advanced Technology Genomics Core (ATGC) for sequencing facility (P30 NIH CA016672).

## Author contributions

T.H.- performed experiments and manuscript writing; M.D.B.- exome-seq data analysis; B.L.- RNA-seq and exome-seq data analysis; M.A.- exome-seq data analysis and intellectual input; M.C.- provided Vk*MYC mice and intellectual input; C.M.A.- conceptualized the study, experimental designing, data analysis, manuscript writing, correspondence

