## Supplementary Table S1 for "B-cell-Specific *Wwox* Deletion Promotes Plasmablastic Tumor Development and Pro-Inflammatory Signatures in a Myeloma Mouse Model"

**Table S1 | Histopathology and Cd138 immunostaining status across plasmablastic tumors**

| **Genotype** | **Mouse ID** | **Histopathology** | **Cd138** | **Cd19** |
| --- | --- | --- | --- | --- |
| *Vk*Myc:Wwox KO* | 101 | Plasmablastic Plasmacytoma | **+++** | **-** |
|  | 117 | B cell lymphoma | **-** | **+++** |
|  | 118* | Plasmablastic Lymphoma | **+** | **+++** |
|  | 170 | B cell lymphoma | **-** | **+++** |
|  | 193 | Plasmablastic Plasmacytoma | **+++** | **-** |
|  | 195* | Plasmablastic Plasmacytoma | **+++** | **-** |
|  | 196 | Plasmablastic Lymphoma | **+++** | **+** |
|  | 197 | Plasmablastic Lymphoma | **+** | **+++** |
|  | 205 | Plasmablastic Plasmacytoma | **+** | **-** |
|  | 206 | Plasmablastic Plasmacytoma | **++** | **-** |
|  | 210 | B cell lymphoma | **-** | **+++** |
|  | 223 | Plasmablastic Lymphoma | **+** | **+++** |
|  | 235* | Plasmablastic Plasmacytoma | **++** | **-** |
|  | 236* | Plasmablastic Plasmacytoma | **+++** | **-** |
|  | 260 | Plasmablastic Plasmacytoma | **+++** | **-** |
|  | 297 | Plasmablastic Lymphoma | **+++** | **+++** |
|  | 298 | Plasmablastic Plasmacytoma | **+** | **-** |
| *Vk*Myc:Wwox WT* | 107 | Plasmablastic Lymphoma | **++** | **++** |
|  | 110 | Plasmablastic Plasmacytoma | **++** | **-** |
|  | 123 | Plasmablastic Plasmacytoma | **+++** | **-** |
|  | 237 | Plasmablastic Plasmacytoma | **++** | **-** |

** Tumor tissues used for RNA-seq and exome-seq analysis*
