## Supplementary Figures for "B-cell-Specific *Wwox* Deletion Promotes Plasmablastic Tumor Development and Pro-Inflammatory Signatures in a Myeloma Mouse Model"

**Supplementary Figure S1**

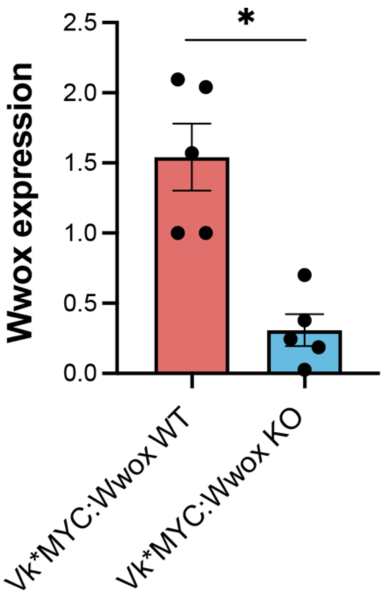


**Supplementary Figure S1 | *Wwox* mRNA expression in Cd19+ B cells.**

Bar graphs illustrate the average mRNA expression levels of *Wwox* determined by qRT-PCR, in Cd19+ B cells purified from the bone marrow of *Vk*MYC:Wwox WT* and *Vk*MYC:Wwox KO* mice (n=5 mice/group). Each data point reflects the expression value for an individual mouse. Error bars indicate the mean ± SEM. The p-value for the loss of *Wwox* expression is $< 0.05$, as determined by an unpaired Student's t-test.

**Supplementary Figure S2**

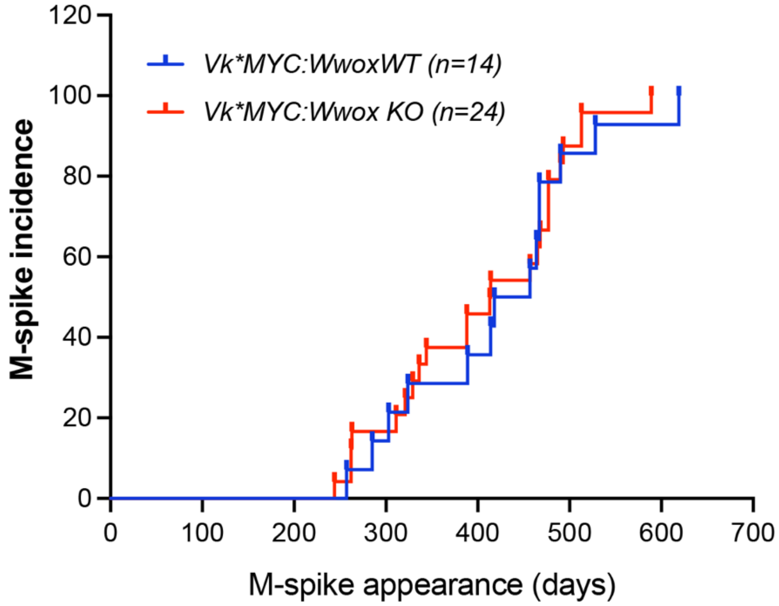


**Supplementary Figure S2 | Age of M-spikes appearance does not differ between *Vk*MYC:Wwox KO* and *WT* mice.**

Kaplan-Meier plot comparing mouse age for M-spikes appearance in and *Vk*MYC:Wwox KO* (red line, n=24) and *Vk*MYC:Wwox WT* (blue line, n=14) mice. The mean age of M-spike onset was 404 days in the *Vk*MYC:Wwox KO* group, with the earliest occurrence at 244 days, compared to 420 days in the *Vk*MYC:Wwox WT* group, with the earliest occurrence at 257 days. The difference between the groups was not significant (Log-rank [Mantel-Cox] p-value = 0.7064).

**Supplementary Figure S3**

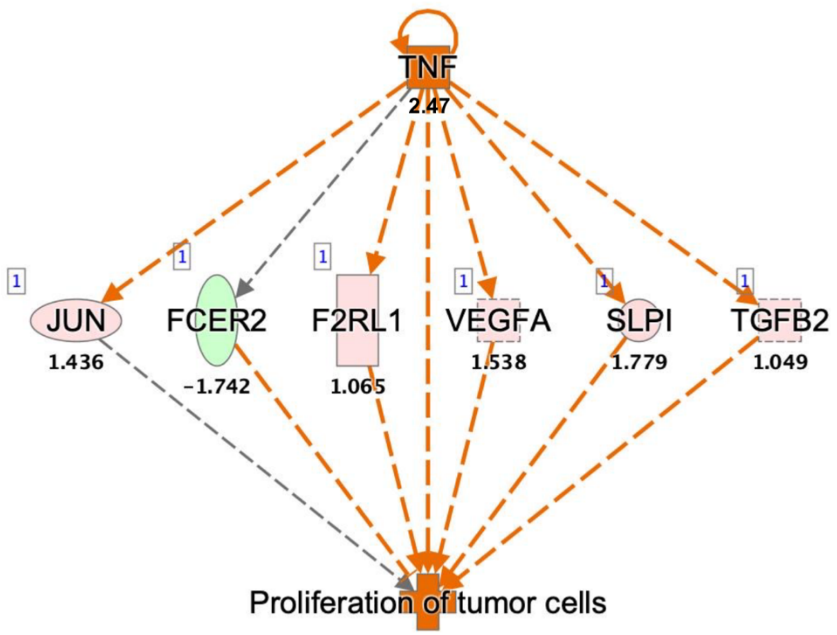


**Supplementary Figure S3 | Tnf-based regulatory effect in *Vk*MYC:Wwox KO* BM cells.**
The topmost activated regulator, Tnf, influenced the regulation of DEGs shown in the network, highlighting 'proliferation of tumor cells' as a significant regulatory effect in *Vk*MYC:Wwox KO* Cd138+ BM cells. The number below Tnf represents the activation Z-score (2.47). The color of the DEG bubbles indicate the direction of regulation (red for upregulated, green for downregulated), and the numbers below DEGs represent gene expression fold change.

**Supplementary Figure S4**

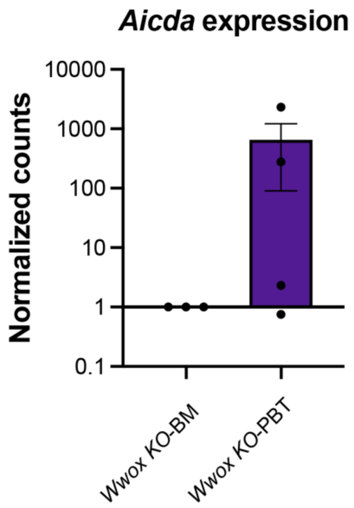


**Supplementary Figure S4 | *Aicda* expression in *Vk*MYC:Wwox KO* plasmablastic plasmacytomas.**

Bar graph illustrates the average normalized counts of *Aicda* mRNA expression in *Vk*MYC:Wwox KO* BM (n=3) and *Vk*MYC:Wwox KO* PBPs (n=4). Each data point represents the expression level for an individual mouse, with error bars indicating the mean ± SEM.

**Supplementary Figure S5**

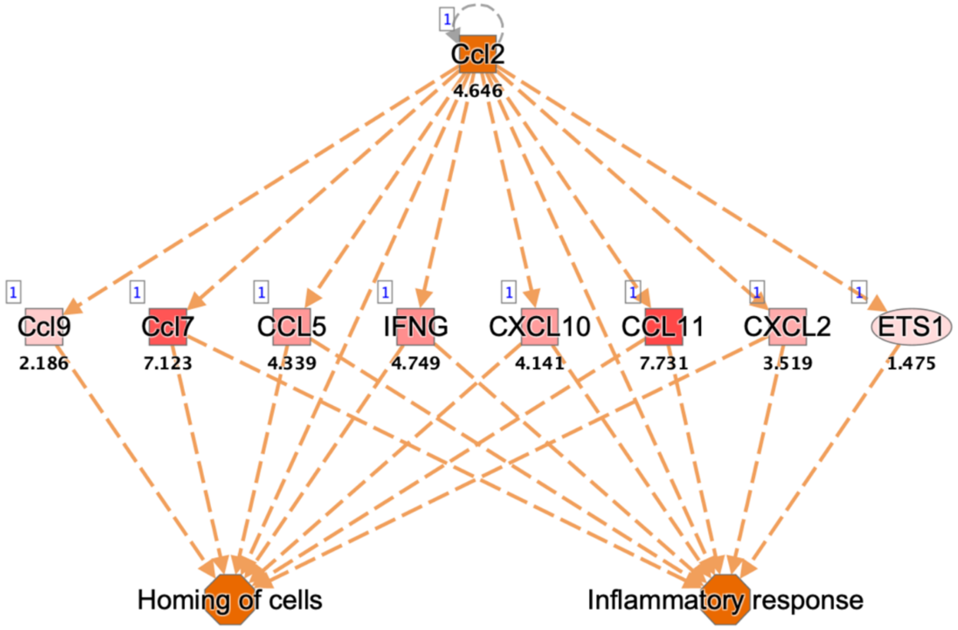


**Supplementary Figure S5 | Ccl2-based regulatory effect in *Vk*MYC:Wwox KO* PBPs.**

The upstream regulator Ccl2 influenced the regulation of multiple target genes shown in the network, highlighting 'Homing of cells' and 'Inflammatory response' as significant regulatory effects in *Vk*MYC:Wwox KO* PBP. The number below Ccl2 represents the activation Z-score (4.64). Red-colored DEG bubbles indicate gene upregulation, with the numbers below representing gene expression fold change.

**Supplementary Figure S6**

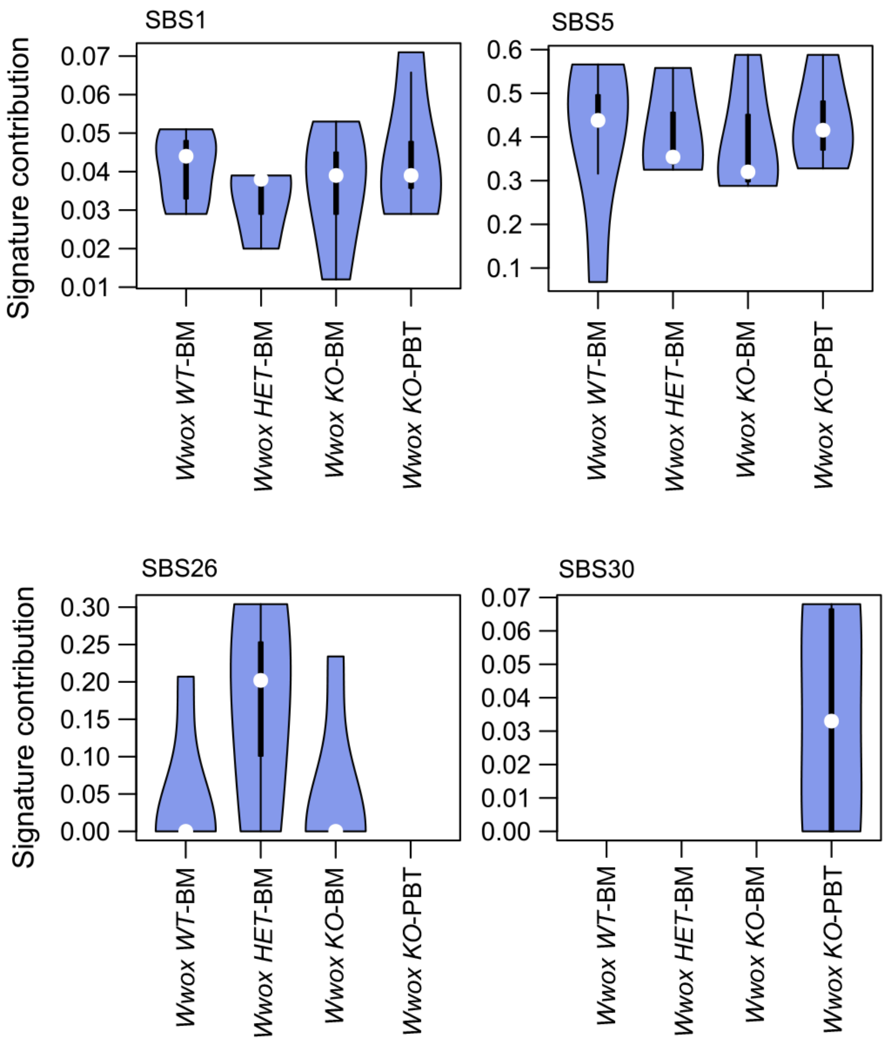


**Supplementary Figure S6 | SBS signatures in BM plasma cells and *Vk*MYC:Wwox KO* PBPs.** Violin plots showing the average contribution values of SBS signatures in BM samples and plasmablastic tumors. The plots represent the signature contributions for SBS1, SBS5, SBS26, and SBS30, as identified in BM plasma cells and *Vk*MYC:Wwox* KO PBPs

**Supplementary Figure S7**

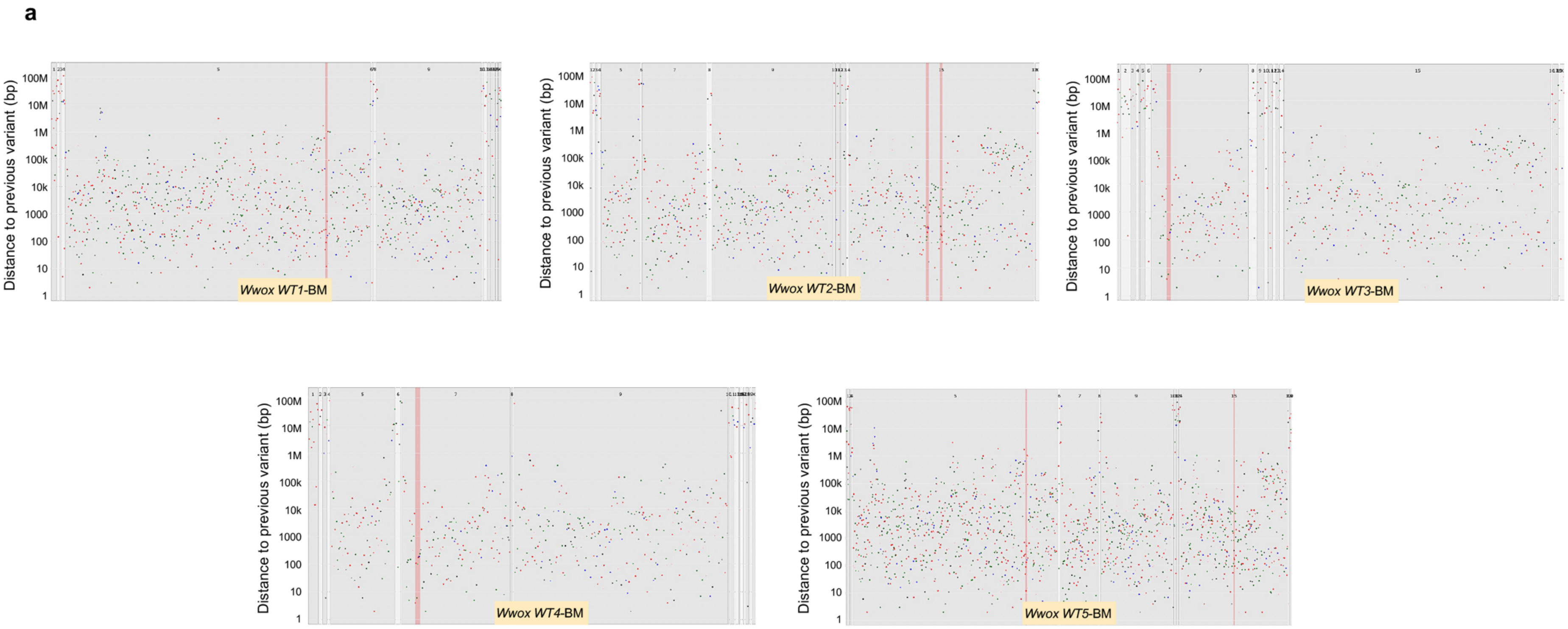

#### Supplementary Figure S7

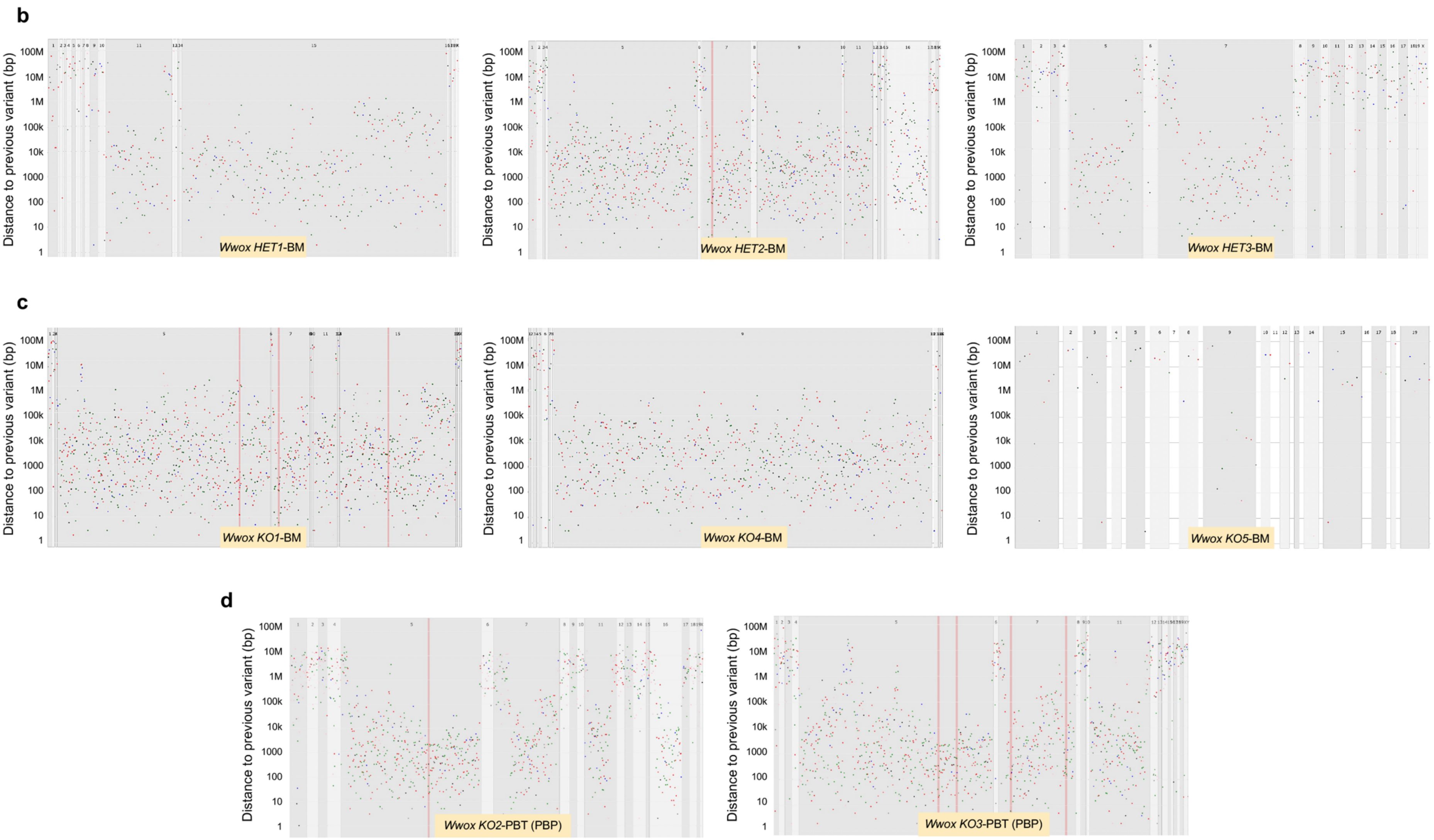

**Supplementary Figure S7 | Mutation profiles in BM plasma cells and *Vk*MYC:Wwox KO* PBPs.**

Rainfall plots displaying the distribution of mutations across the genomes of BM plasma cell samples and *Vk*MYC:Wwox* KO PBPs. The Y-axis represents the distance of each mutation from the preceding mutation, while different colors indicate the various types of mutation substitutions.

**Supplementary Figure S8**

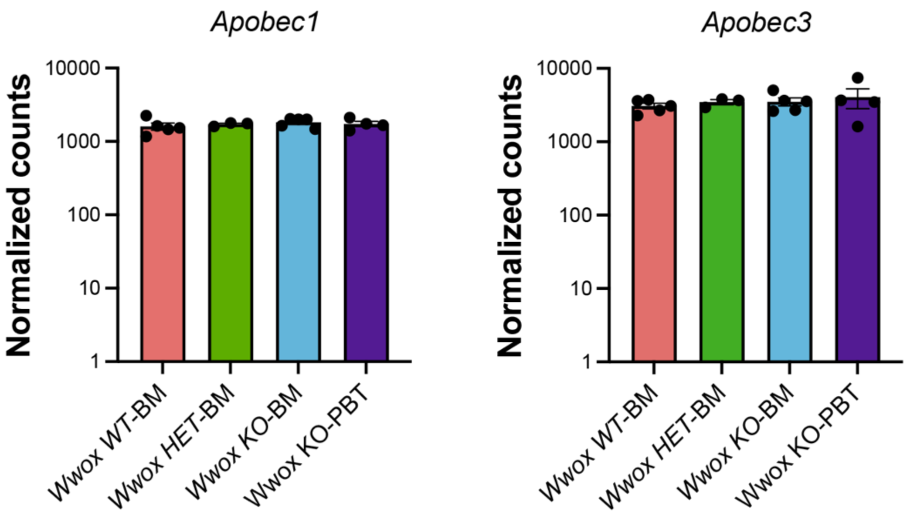


**Supplementary Figure S8 | Apobec1 and Apobec3 expression in BM plasma cells and *Vk*MYC:Wwox KO* PBPs.**

Bar graphs illustrating the average normalized counts of *Apobec1* and *Apobec3* mRNA expression in *Vk*MYC:Wwox WT* (n=5), *HET* (n=3), *KO* (n=5) BM and *Vk*MYC:Wwox KO* PBP (n=4) samples. Each data point represents the expression level for an individual mouse, with error bars indicating the mean ± SEM.

### Supplementary Figure S9

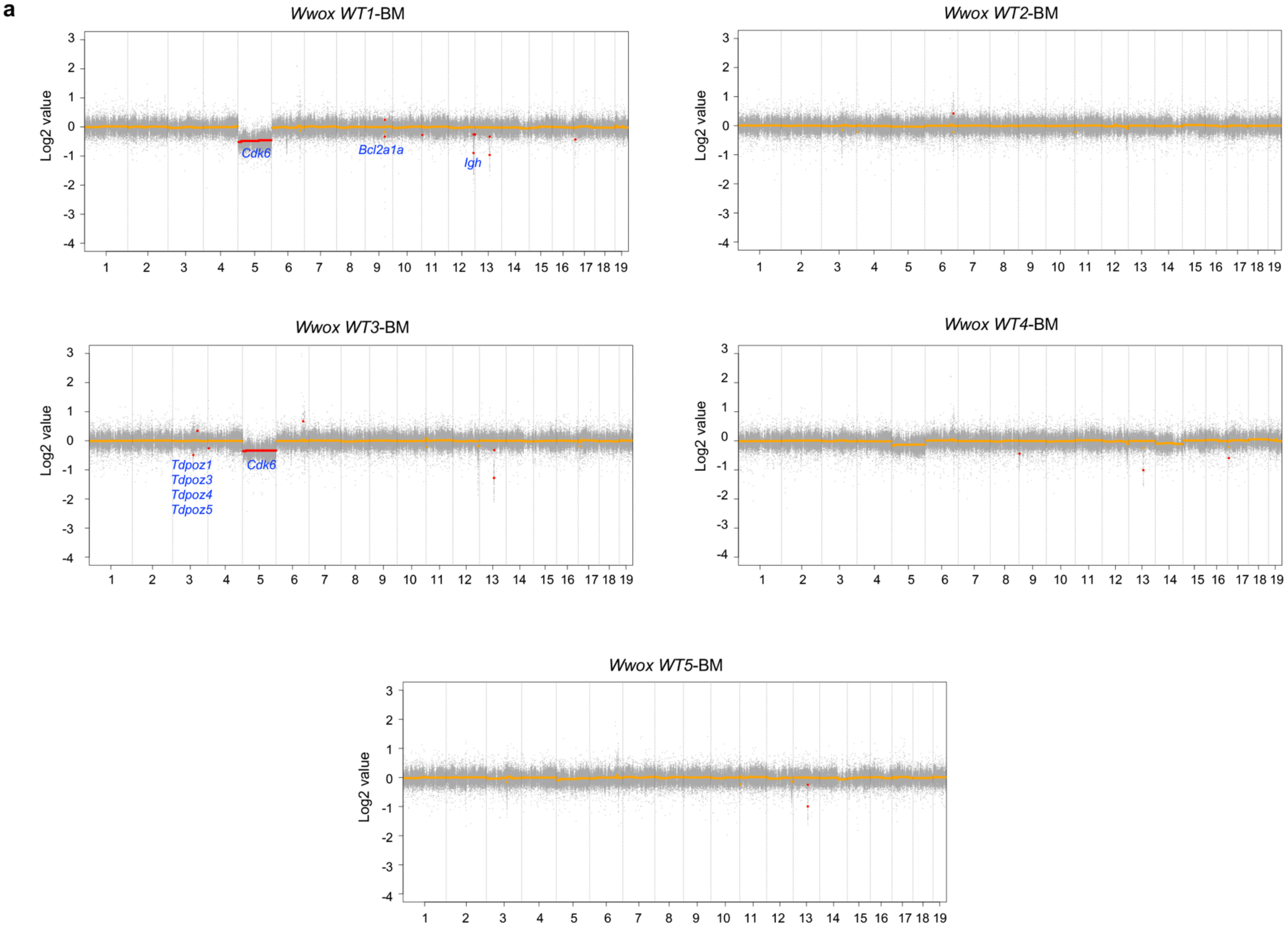

#### Supplementary Figure S9

**b**

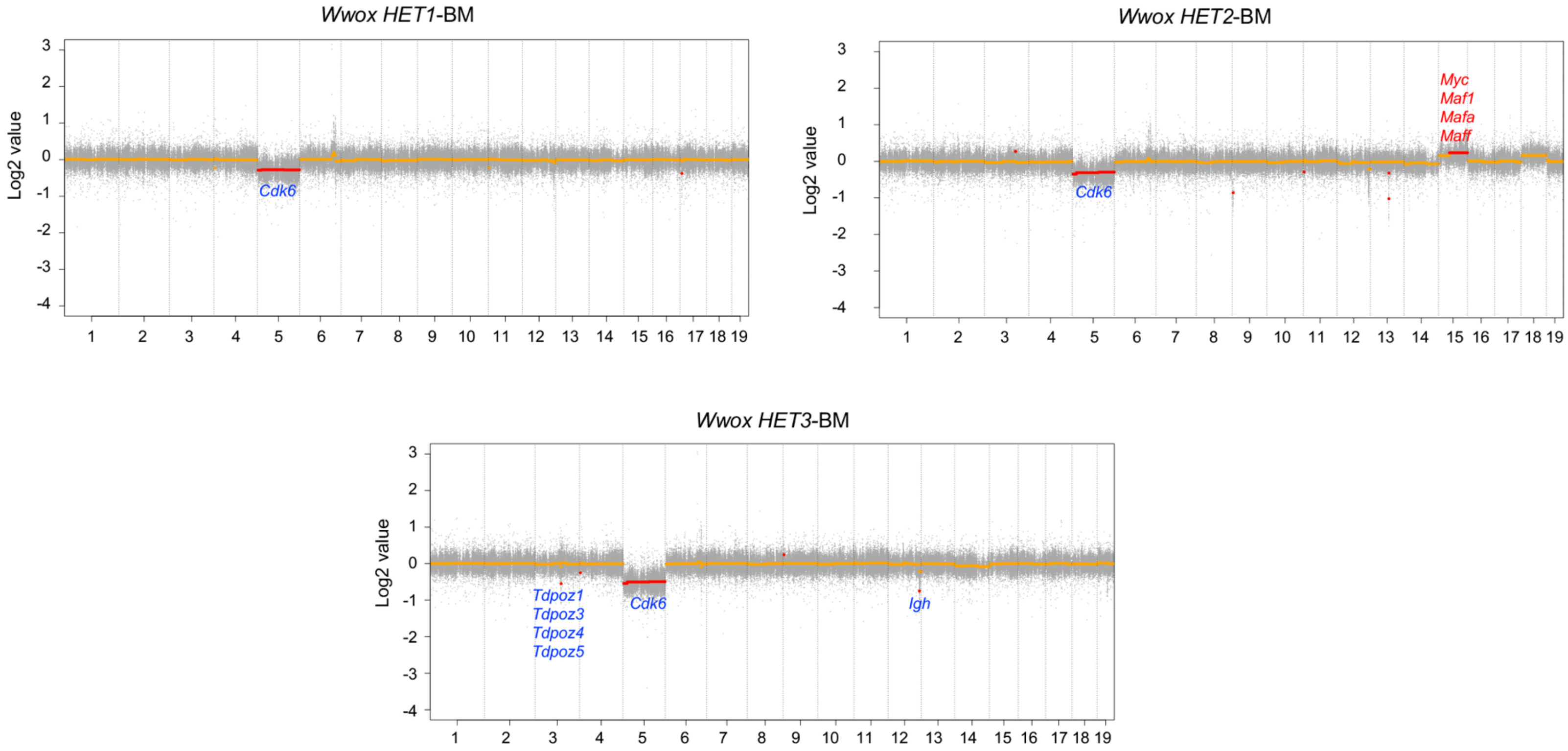

**Supplementary Figure S9**

**c**

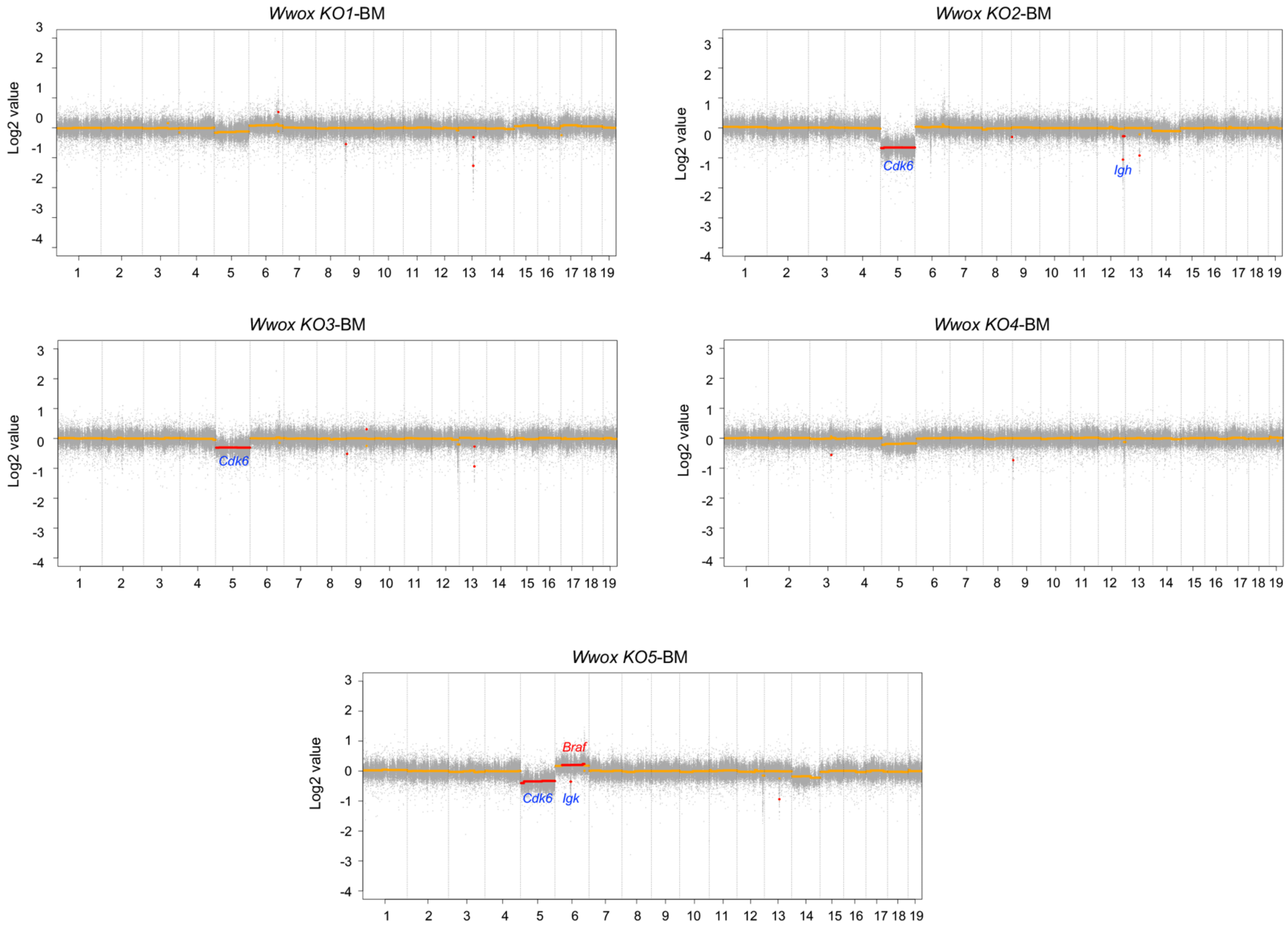

**Supplementary Figure S9 | Focal and large copy number alterations in BM plasma cells and *Vk*MYC Wwox KO* PBPs.**

Scatter plots from BM plasma cell samples and Vk*MYC Wwox KO PBPs. Focal and large copy number alterations are represented by red markers; those above zero indicate copy number gains, while markers below zero signify copy number losses. The names of the genes affected by specific copy number alterations are displayed near the corresponding alterations, with red indicating copy number gains and blue indicating copy number losses for the respective genes.
